# Early life milk diets shape infant gut microbiota: evidence of microbial plasticity in response to breast and formula milk

**DOI:** 10.1101/2024.09.22.614312

**Authors:** Melissa AE Lawson, Matthew Dalby, Shabhonam Caim, Larissa Richardson, Albert Koulman, Raymond Kiu, Gwen LeGall, Lindsay J Hall

**Affiliations:** Food, Microbiome and Health, Quadram Institute Bioscience, Norwich Research Park, Norwich, UK; Lydia Becker Institute for Immunology and Inflammation; Wellcome Trust Centre for Cell Matrix Research; Division of Infection, Immunity and Respiratory Medicine; School of Biological Sciences, Faculty of Biology, Medicine and Health, University of Manchester, Manchester Academic Health Science Centre, Manchester, UK; Department of Microbes, Infection and Microbiomes, College of Medicine and Health, University of Birmingham, Birmingham, UK; Institute of Microbiology and Infection, University of Birmingham, Birmingham; Wellcome-MRC Institute of Metabolic Science, University of Cambridge, Cambridge, UK; Norwich Medical School, University of East Anglia, Norwich Research Park, Norwich, UK

**Keywords:** Microbiota, early life, breast milk, formula, lipids, metabolites

## Abstract

In early life, diet plays a key role in shaping the infant microbiota, yet the impact of breast milk and formula on microbial ecosystems at different stages of infant development remains poorly understood. Here, we performed static batch culture experiments using infant faecal samples at ages 1, 12 and 18 months of age, supplemented with either breast milk or formula for 48 hours. We further removed small metabolites (e.g. small carbohydrates, proteins and lipids) from breast and formula milk through dialysis and also added this to the static batch cultures with faecal samples from each infant. Our results show that the one-month-old faecal microbiota exhibited the greatest sensitivity to dietary intervention, with significant changes in microbial composition, metabolites and lipids, particularly in response to formula supplementation. In contrast, the microbiota at 12 months displayed increased stability, while the 18-month-old infant samples were the most resilient to different dietary supplementation. We also observed that triglyceride (TG46:1) was produced in the youngest infant samples but consumed by the older infant microbiota, suggesting a shift in metabolic interactions as the gut microbiota diversifies with age. Metabolite profiles linked to KEGG pathways further indicated the older infants diverse microbiotas had greater functional capacity in comparison to the one-month-old infant, particularly in response to breast milk. Fewer overall metabolic pathways were affected when the infant samples were grown in formula. Collectively, our data underscores the importance of early life diet in shaping microbiota profiles, with more stable microbial ‘dynamics’ in more mature gut ecosystems, in response to dietary changes which are also associated with downstream functional (i.e. metabolite/lipid) readouts. Notably, the removal of small metabolites from breast milk by dialysis highlighted the potential role of milk lipids in promoting growth of foundational microbiota members like *Bifidobacterium*. Our data, along with further studies, are required to probe the mechanisms by which specific nutrients, particularly lipids, modulate microbial composition and support infant health during the first 2 years of life.

## Introduction

The early life microbiota of a healthy infant is typically established through vertical transmission from the mother’s microbiome (Ferretti et al., 2018; Yassour et al 2018). Initially, the infant gut microbiome is low in species diversity and as the infant grows, the gut microbiome evolves in complexity and diversity that is influenced by various external factors, with diet being the most significant determinant (Laursen et al., 2017). The most dramatic change in microbial diversity occurs during the first few years of life, as this coincides with the microbiota maturing from a newborn to toddler stage (Stewart et al., 2018; Laursen et al., 2017). Infants exclusively fed maternal breast milk for the first six months typically exhibit a microbiota characterised by low diversity microbiota but enriched with beneficial Gram-positive bacteria like *Bifidobacterium* (Ma et al., 2020). Breast milk is nutritionally optimised to support infant growth, and as such is recommended as the exclusive source of nutrition for their first six months of life by global health organisations (UNICEF-WHO 2018). In addition, breast milk contains unique components such as human milk oligosaccharides (HMOs), which selectively enhance the growth of key microbiota members, including certain *Bifidobacterium* species and strains (Laursen et al., 2021; Elson et al., 2016). The presence of *Bifidobacterium* in the infant gut has been linked to key health outcomes, including immune system modulation, reduced risk of autoimmune and allergic diseases, and production of essential nutrients and vitamins that promote overall child well-being (Sun et al., 2021; O’Neill et al., 2017; Björkstén et al., 2001); thereby making *Bifidobacterium* a critical component of the early life gut microbiota.

The link between exclusive breast-feeding and high levels of *Bifidobacterium* levels in the infant gut has been extensively documented (Lyons et al., 2022; Ma et al., 2020; Stewart et al., 2018). Moreover, *Bifidobacterium* plays a vital role in digestion of breast milk, liberating key milk-specific nutrients such as monosaccharides and oligosaccharides, that can support the growth of other bacterial species within the infant gut microbiota (Lawson and O’Neill 2020; De Vuyst and Leroy 2011). Breast feeding may also seed the infant gut with *Bifidobacterium*, further highlighting the relationship between infant diet and microbiota composition (Lyons et al., 2022; Laursen et al., 2021; Pannaraj et al. 2017). It is clear that a simple but stable gut microbiota characterises the gut microbiome of solely breastfed infants, with minimal changes occurring unless the diet changes. As infants grow, their microbiota undergoes dynamic evolution driven by dietary transitions. Dietary transitions can occur internally, by tchanges in mother’s breast milk composition in response to her diet and developmental cues from the infant (i.e. milk demand); while externally, a major transition occurs when solid foods are typically introduced around 6 months of age (Cheema et al., 2022; Lyons et al., 2022). These dietary shifts are critical in shaping the composition and function of the microbiota during this formative period, and the change continues for the first few years of life.

Whilst the benefits of breastfeeding are well established (UNCEIF-WHO, 2018), not all parents are able to or choose to exclusively breast-feed and many reply on formula milk as an alternative. In fact, there are many mothers that have self-reported to use formula (either alone or in combination with breast milk) (Victora et al., 2016; Rollins et al., 2016). Supplementation with formula is associated with significant changes in the diversity and composition of the gut microbiota compared to exclusive breastfeeding (Ma et al., 2020). Despite the efforts of manufacturers to replicate breast milk, the composition of infant formulas can vary widely based on manufacture, region and even batch (Furse and Koulman 2019). Several studies have demonstrated differences in microbial composition between breastfed or formula-fed infants, but it remains unclear how early or late in development the microbiota becomes stable and if the infant gut microbiome has a memory response upon secondary exposure to a milk diet. Interestingly, evidence from studies on HMOs supplemented adults indicates that diet-induced shifts (or nutritional microbial memory) may extend beyond infancy, with long-term impacts on the microbiome (Elison et al., 2016). In this study, we explored how the infant gut microbiota responds to different milk diets (breast milk and formula) at three key developmental stages (1, 12 and 18 months) in static batch cultures. Using stool samples from healthy infants, we examined changes in microbiota composition, metabolite production and lipid profiles in response to the sole energy source being either breast or formula milk. Our findings highlight a high degree of plasticity in the infant gut microbiota during the first few months of life, with dramatic shifts in microbial composition observed in the one-month-old infant in response to diet. In contrast, the microbiota of older infants (12 and 18 months) displayed greater stability, with the eldest infant showing a form of ‘microbial memory’, reverting to a simpler microbiota dominated by early life bacteria, such as *Bifidobacterium* when grown in breast milk.

## Methods

### Infant faecal samples

Stool samples were collected from three full-term, healthy infants at the age of (one month, 12 months and 18 months), and were processed within 24h. None of the infants had received antibiotics within the one-month prior to sample collection. The one-month-old received a sole milk diet (2 weeks breast milk; 2 weeks formula milk); the 12-month old’s diet was predominantly breast milk until 11 months (with simple solid foods) and had been transitioned to solid foods during the day given a bottle of breast milk each evening. The oldest infant was breast-fed, and at the time of collection was eating a diet of simple and complex foods, with a bottle of formula prior to bed. At the time of collection none of the infants had received antibiotics/probiotics prior to sampling, and collection was in accordance with protocols laid out by the National Research Ethics Service (NRES) approved UEA/QIB Biorepository (Licence no: 11208) and Quadram Institute Bioscience Ethics Committee (see Table S1 for infant metadata).

### Breast and formula milk

Donated expressed breast milk was collected at home and frozen from four different mothers, over their child’s first year of life. All milk was tested for the growth of Bifidobacterium on de Man Rogosa and Sharpe (MRS) media with mupirocin and l-cysteine (0.05 mg/mL each, Sigma-Aldrich, Dorset, UK). Undergoing a single thaw, all the milk was pooled, and half of the volume of the pooled breast milk underwent dialysis in using SnakeSkin™ Dialysis Tubing, 3.5K MWCO (Sigma) in dialysis buffer (0.05M NaH_2_PO_4_ H_2_O, 0.3M NaCl, pH 8.0; changed every 12 hours) for 2 days at 4°C, while the remainder was stored at 4°C for 2 days. All milk (both whole and dialysed milk) was freeze dried on day 3, then further mixed and stored in an airtight container until use (temp: -47°F, vacuum: 117-121 mBar; Modulyo®D Freeze Dryer, Thermo). Formula milk was first hydrated according to manufacturer’s directions (Aptimal® First milk), mixed between batches bought and then prepared in parallel with the breast milk.

### Culture conditions

Infant stool was homogenised in sterile PBS and the equivalent of 0.5g of faecal slurry was added to minimal media (Sigma) supplemented with 20% w/v of milk and grown in an anaerobic chamber at 37°C (Don Whitley Scientific, Bingley, UK; containing 5% CO_2_, 10% Hydrogen, 85% Nitrogen gas). Samples were taken every hour from 0h to 10h, 24h, 30h and 48h post inoculation.

### 16S rRNA amplicon library preparation and bioinformatics analysis

The FastDNA Spin Kit for Soil (MPBIO, California, USA) was used DNA extraction according to manufacturer’s protocol, with the following exception of two additional (30s) bead-beating steps prior to extraction. The 16S rRNA gene was amplified using primers binding to the V1-V2 16S rRNA gene as described in (Alcon-Giner et al. 2017). Quality control of Illumina MiSeq Raw reads was performed using FASTX-Toolkit53 with a minimum quality threshold of 33 for at least 50% of the bases. All passed read (separately for both pairs) were aligned against the SILVA database (Quast et al. 2013) using BLASTN55 (Altschul et al. 1997), and then annotated using the paired-end protocol in MEGAN (Huson, Mitra, and Ruscheweyh 2011). PCA plots were generated with Eigen vector normalisation. All raw sequence reads are now stored under project accession PRJNA1161489 at NCBI.

### ^1^H NMR

Cultured samples were thawed at room temperature, filtered to remove large lipid/proteins (that block ^1^H NMR signals). The remaining solution was then centrifuged at max speed for 10 min, and 50mg of sample was mixed with 600 μL NMR buffer (1 mM TSP) in preparation for ^1^H NMR spectroscopy. Each ^1^H NMR spectrum was acquired with 512 scans, a spectral width of 12300 Hz and had an acquisition time of 2.7 s. The “noesypr1d” presaturation sequence was used to suppress the residual water signal with a low-power selective irradiation at the water frequency during the recycle delay. Spectra were transformed with a 0.3-Hz line broadening, manually phased, baseline corrected and referenced by setting the TSP methyl signal to 0 ppm. Metabolite identification was based on the previously published databases (1-5), including the web (Human Metabolome Database, http://www.hmdb.ca/) and using the 2D-NMR methods, COSY, HSQC, and HMBC. Metabolite quantification was assessed using Chenomx NMR suite 7.6™ software. KEGG Pathway assignment was analysed using MetaboAnalyst version 3.0 (McGill University).

### Lipidomics sample extraction

150μL was removed from each flask and snap-frozen in liquid nitrogen, prior to lipid profiling. An automated method for the extraction of lipids was developed using an Anachem Flexus automated liquid handler (Anachem, Milton Keynes, UK). 100 μL of MilliQ H2O was added to each of the wells and mixed, and then 100 μL of the mixture was transferred to a glass coated 2.4 mL deep well plate (Plate+™, Esslab, Hadleigh, UK). Next, 250 μL of MeOH was added containing 16 internal standards (CE(18:0-d6); Ceramide C16-d31; free fatty acids: C15:0-d29, C17:0-d33, C20:0-d39; LysoPC(C14:0)-d42, PA(C16:0-d31/C18:1); PC(C16:0-d31/C18:1); PE(C16:0-d31/C18:1); PG(C16:0-d31/C18:1); PI(C16:0-d31/C18:1); PS(C16:0-d62); SM(C16:0-d31); TG(45:0-d87); TG(48:0-d93); TG(54:0-d105)), followed by 500 μL of methyl tert-butyl ether (MTBE). The plates were then sealed (using Corning aluminium micro-plate sealing tape [Sigma Aldrich Company, UK]) and shaken for 10 min at 600 rpm, after which the plate was transferred to a centrifuge and spun for 10 min at 6000 rpm. Each well in the resulting plate had two layers, with an aqueous layer at the bottom and an organic layer on top. A 96-head micro-dispenser (Hydra Matrix, Thermo Fisher Ltd, Hemel Hampstead, UK) was used to transfer 25 μL of the organic layer to a glass coated 240 μL low well plate (Plate+™, Esslab, Hadleigh, UK), and 90 μL of MS-mix (7.5 mM NH4Ac IPA:MeOH [2:1]) was added using a Hydra Matrix, after which the plate was sealed and stored at –20°C until analysis.

Direct infusion high-resolution mass spectrometry. All samples were infused into an Exactive Orbitrap (Thermo, Hemel Hampstead, UK), using a Triversa Nanomate (Advion, Ithaca, US). The Nanomate infusion mandrel was used to pierce the seal of each well before analysis, after which, with a fresh tip, 5 μL of sample was aspirated, followed by an air gap (1.5 μL). The tip was pressed against a fresh nozzle and the sample was dispensed using 0.2 psi nitrogen pressure. Ionisation was achieved with a 1.2 kV voltage. The Exactive started acquiring data 20 seconds after sample aspiration began. The Exactive acquired data with a scan rate of 1 Hz (resulting in a mass resolution of 65,000 full width at half maximum [fwhm] at 400 m/z). After 72 seconds of acquisition in positive mode the Nanomate and the Exactive switched over to negative mode, decreasing the voltage to -1.5 kV. The spray was maintained for another 66 seconds, after which the analysis was stopped and the tip discarded, before the analysis of the next sample began. The sample plate was kept at 15°C throughout the analysis. Samples were run in row order and repeated multiple times if necessary to ensure accuracy.

Lipidomics data processing and High Resolution Mass Spectrometry Peak picking. The data were output from the Exactive in a compressed proprietary raw format. Files were decompressed and converted to mzXML format using the “msconvert” tool in ProteoWizard.20,21 For each infusion an average spectrum is calculated from the user defined time window. Using a mass-to-charge ratio (m/z) window of 185–1000 and a retention time window of 20–70 seconds for positive ionisation mode and 95–145 seconds for negative ionisation mode, the R package “xcms”3 was used to average fifty spectra per mode. The intensity of peaks was extracted using in-house algorithm that used a list of target m/z for lipids expected in these samples (for these samples those are triglycerides. The intensity of the found lipids was expressed relative to the internal standards.

## Results

We employed a multi-omics approach to examine how gut (faecal) microbiota of three infants at different ages (1, 12 and 18 months respectively, Fig S1) responded to breast milk or formula under static experimental conditions; using a combination of 16S rRNA amplicon profiling, lipidomics and metabolomics analysis (Fig 1A). This approach allowed us to explore microbial composition as well as metabolic and lipid changes triggered by microbiota within each milk type.

**Figure 1:**
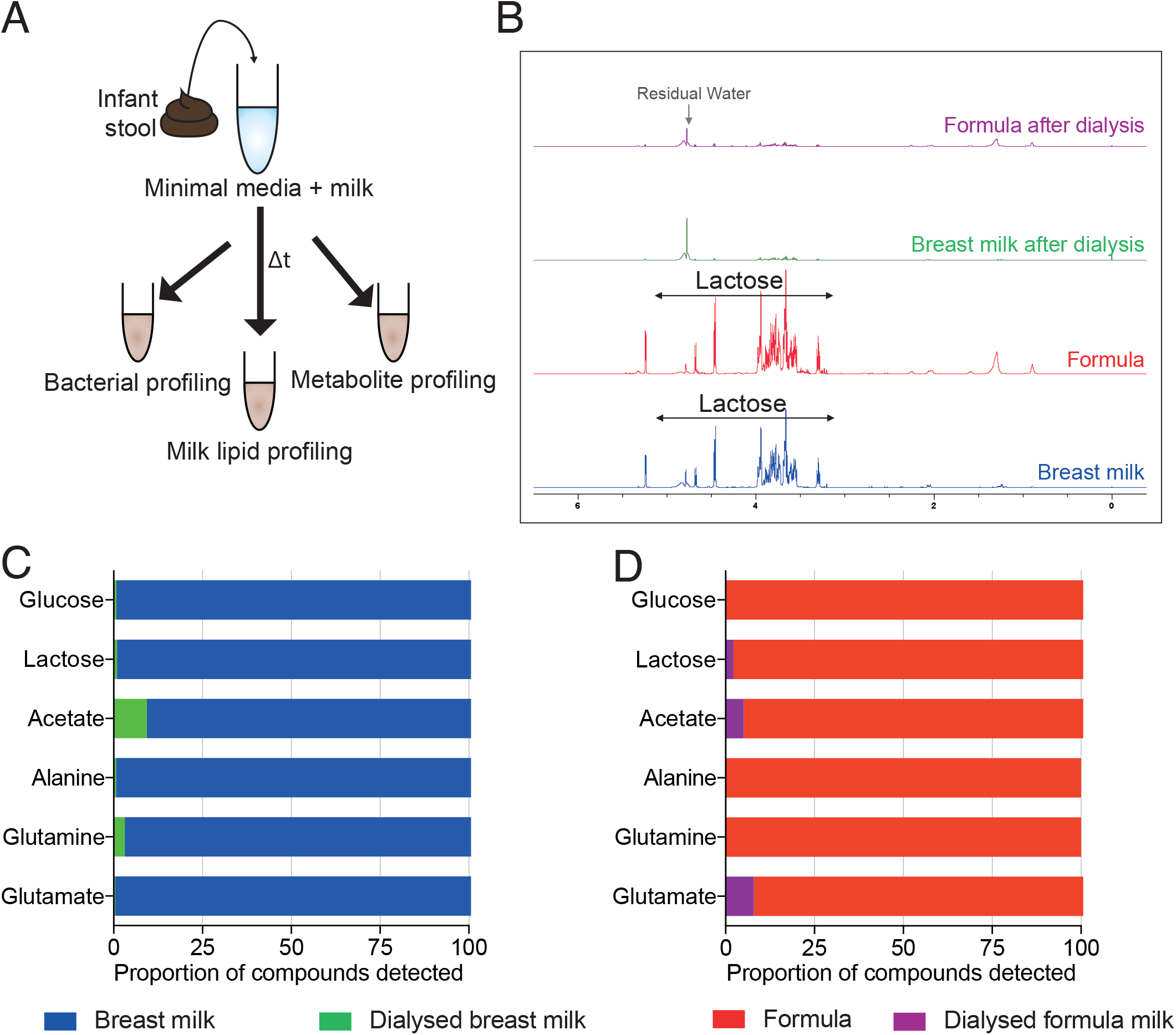
Experimental design to evaluate how the gut microbiota of infants at one, 12, and 18 months of age change in response to a milk diet. (A) Schematic illustrating experimental set-up and sampling procedure. (B) ^1^H-NMR analysis indicating the metabolite profile of breast and formula milk before and after dialysis. The most abundant sugar in milk, lactose has been indicated on the plots. (C) The proportion of sugars, organic and amino acids present before and after dialysis (a full list of compounds are available in Table S1).

Both breast and formula milk provide a variety of complex nutrients to the infant and their developing gut microbiome. As the infant consumes milk, it passes through the gastrointestinal tract where key nutrients are absorbed. After entering the stomach, milk slurries move into the intestines, where many simple sugars (like glucose and lactose), and other small metabolites are largely absorbed by host epithelial cells in the upper small intestine. By the time milk slurries reaches the large intestine, where the densest collection of gut microbes resides, most simple sugars and readily absorptive compounds have already been depleted (Table S2). Therefore, to emulate the exposure of gut bacteria like *Bifidobacterium* to milk in the infant colon, we processed the pooled milk samples by dialysis to remove smaller metabolites and compounds. Both breast and formula milk were dialysed to generate slurries that were depleted of simple sugars, small peptides and metabolites (≤ 3.5KDa in size) including, but not limited to, lactose, glucose, smaller HMOs (including 2’FL and LNT), and amino acids including alanine, glutamine, glutamate (Figure 1B-D, Table S2). This remaining milk slurry we used in experiments contained the larger components of milk-like casein and whey proteins, and larger lipid droplets, in an attempt to closely mimic what the colonic microbiota would likely be exposed too. Throughout the study, the dialysed breast milk and dialysed formula slurries were compared to their undialysed counterparts (referred to as breast and formula milk) to evaluate the microbial, lipid and metabolic responses in each condition.

### Diet-induced microbial changes

We first characterised the faecal microbiota profiles of the three infants prior to the start of the experiment and found that each infant had a unique faecal microbial composition, which, as expected increased in diversity with age (Fig S1). These infant faecal samples were then used to inoculate our batch culture systems and microbial and metabolite/lipidomic profiles were monitored over 48 hours in response to the different milk diets. Using 16S rRNA amplicon sequencing to track microbial changes over time, we identified distinct responses to each experimental condition based on the infants age (Fig S2A-B, Figure 2S).

**Figure 2:**
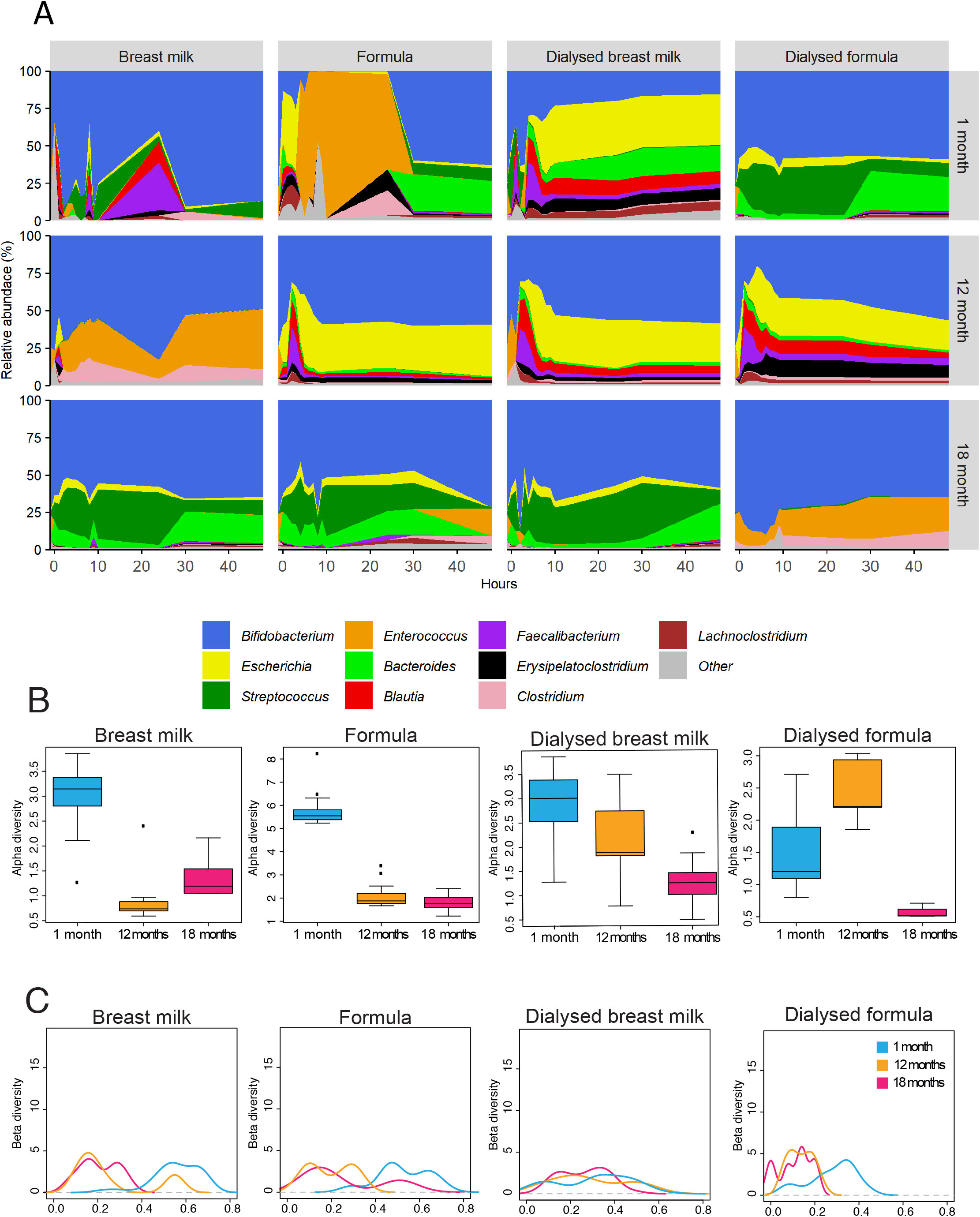
Milk-induced microbial changes over 48h in faecal slurries from a one, 12 and 18-month infant. (A)16S rRNA gene profile of top 10 genera present over time. Changes in the (B) alpha diversity and (C) beta diversity for each microbial profile over 48h growth in minimal media with either whole breast milk, formula, dialysed breast and dialysed formula milk.

#### Whole breast milk

The majority of 16S amplicon reads across all ages belonged to the genus *Bifidobacterium*, (Fig 2A) consistent with existing research that indicates the preferential growth of *Bifidobacterium* spp. in breast-fed infants (Koenig et al., 2011; Yatsunenko et al., 2012; Bäckhed et al., 2015). Surprisingly, the faecal samples from the one-month-old infant had the highest microbial diversity in breast milk compared to the results from the two older infants (Fig 2B-C). In contrast, the 12-month sample showed a composition dominated by *Bifidobacterium, Enterococcus* and *Clostridium*, while the 18-month sample was enriched in *Bifidobacterium, Bacteroides* and *Streptococcus* (Fig 2A). These findings suggest that breast milk, regardless of age consistently drives a shift toward a more simplistic microbiota composition dominated by a small number of genera, regardless of the infant’s age.

#### Whole formula milk

Again, in formula milk, we observed the faecal microbiota of the one-month-old infant had a highly dynamic and stochastic composition, with significant changes observed over the 48-hour period (Fig 2A-B). Early timepoints were dominated by *Escherichia, Blautia, Bacteroides and Lachnoclostridium*, which were later replaced by *Enterococcus*, and finally *Bifidobacterium, Bacteroides* and *Streptococcus* at later timepoints (Fig 2A). The 12-month-old sample maintained an increased diversity throughout the experiment, and then ‘stabilised’ into a microbial community largely dominated by *Bifidobacterium*. This pattern was mirrored in the 18-month-old faecal sample, suggesting that older infants develop a more stable and diverse microbial community when exposed to formula (Fig 2A).

#### Dialysed breast milk

In this condition, we propose that the removal of small molecules from breast milk will preferentially feed the older infants’ more diverse microbiomes (Fig S1). However, we found that the one-month-old sample again had the greatest microbial diversity compared to the older infants (Fig 2A-B). Across all age groups, *Bifidobacterium* remained a prominent genus, but the one-month and 12-month-old samples also had a large proportion of *Escherichia* and *Blautia;* whereas the 18-month sample was characterised by a high proportion of reads assigned to *Bifidobacterium* and *Streptococcus*.

#### Dialysed formula milk

We observed minimal changes in microbial diversity over time, except for a transient increase in diversity in the 12-month-old sample within the first 12 hours (Fig 2A-B). Both the youngest and the oldest infant exhibited low turbidity and poor growth, suggesting that the absence of sugars and other smaller molecules lost during dialysis of formula limited bacterial growth. Collectively, these findings suggest that small metabolites present in formula are critical for supporting microbial growth, and in our study particularly in younger infants.

### Microbial diversity and age-related trends

Beta diversity within each infant’s microbiota increased over time in response to each milk condition (breast milk, formula, dialysed breast milk and dialysed formula) and positively correlated with the age of the infant (Fig 2C). Similar trends in beta diversity were observed between the 12-month- and 18-month-old samples, likely reflecting the similarity in their baseline faecal microbiota profiles (Fig S1A-B). Contrary to our initial expectations, the one-month-old infant exhibited the highest plasticity in microbiome composition across multiple conditions, while the older infants displayed more stable microbiomes. This may be attributed to either the early developmental stage of the one-month-old microbiome, and/or the potential influence of early exposure of formula at 2-weeks of age, which may have accelerated diversification. Additionally, the observed stability in the older infants suggests the presence of a “microbial memory”, where previous exposure to breast milk during early life establishes a resilient *Bifidobacterium*-dominated microbiota that responds predictably to re-exposure to milk diets.

### Lipid profiles change over time, regardless of age

Fat molecules are key constituents of breast milk and play a critical role in nutrient and energy storage in infants. Upwards of 50% of the total energy content in breast milk comes from milk fat, (Kim and Yi, 2020) with triglycerides (TG) making up 95-98% of the fatty acids present, and TG46:1 being the most common. These TGs can be enzymatically broken down in the infant gastrointestinal tract into other nutritional components such as free fatty acids, monoglycerides, diacylglycerides and glycerol. In the whole milk conditions, the infant microbiota has access to these lipids, whilst in the dialysed milk conditions, lipid availability was reduced during the dialysis process. Given that small dietary changes can influence the composition of the gut microbiota, we hypothesised lipid availability, namely TG46:1, might drive some of the changes observed in Fig 2. To explore this, we examined the lipidomic profiles, with emphasis on total TG and monoglycerides/diacylglycerides concentrations in each milk condition for each infant using mass spectrometry (Table S3A-C). We found that dialysis had a much greater impact on the breast milk lipidomic profile compared to formula (Fig 4B, S5 and S6). Approximately 80% of the total lipids were lost from breast milk during dialysis, while only about 50% were lost in formula (Fig 4B, S5 and S6). In the undialysed conditions, we observed rapid uptake of breast milk lipids by the fecal microbiota across all infants, while formula lipids showed little effect (Fig 3A and S4). When grown in breast milk the TG and monoglyceride/diacylglyceride concentrations of all three infants increased over time (Fig 3B & S4). However, the increase in the level of TG in the one-month-old by 48h was >2,000 times higher than their older counterparts (Table S1A-C and Fig 3B). This suggests a differential utilisation of lipids between younger and older infants, where the younger infant microbiomes may utilise milk lipids more for growth and energy storage.

**Figure 3:**
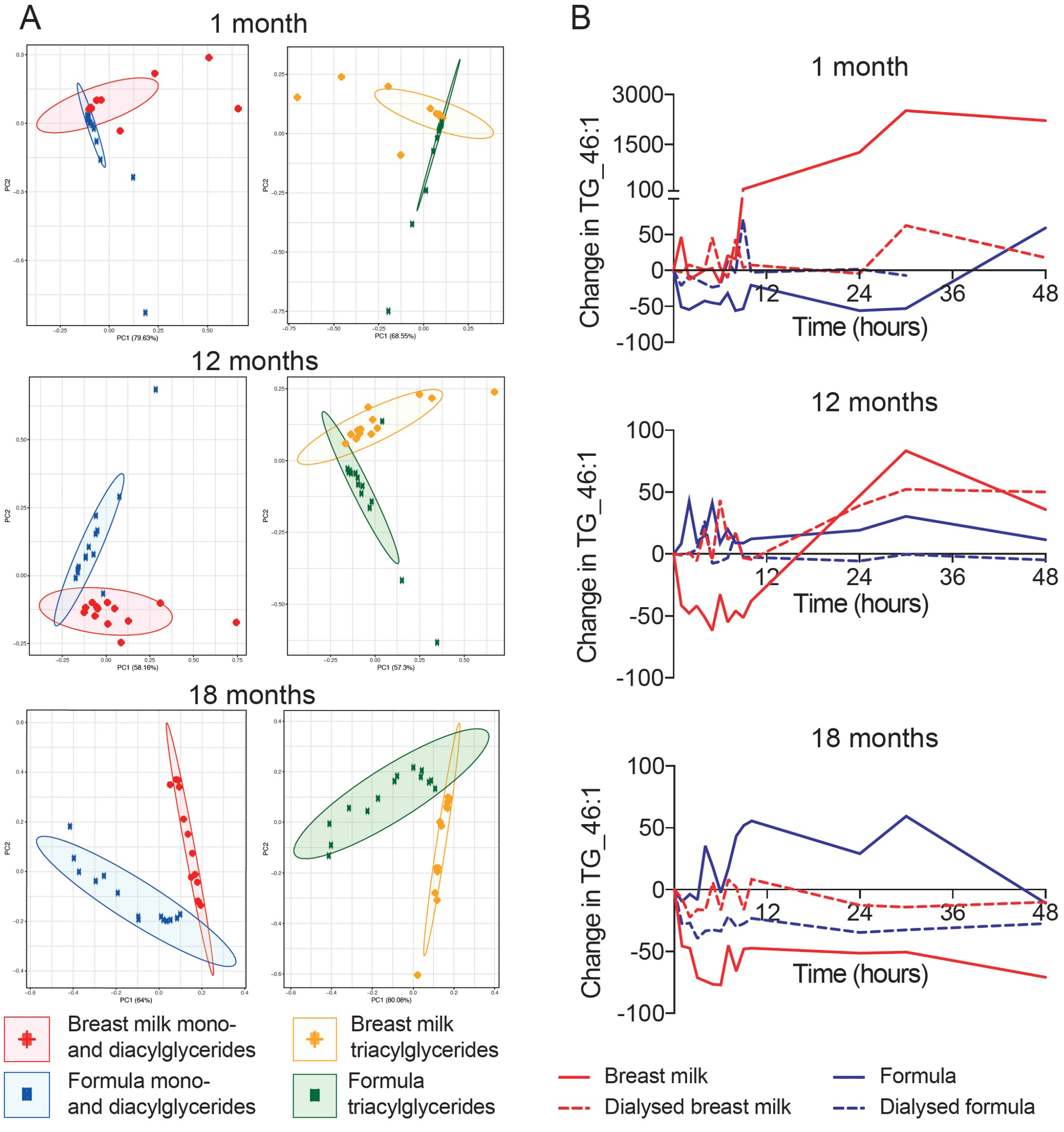
Changes in lipids over time. (A) PCA plots illustrating the differences in the total TG and monoglycerides/diacylglycerides in whole breast milk compared to whole formula for each infant within the first 24h. (B) the absolute change in concentration of TG_46:1 (AU) at each time point measured for each infant faecal sample grown in whole breast milk (solid red line), whole formula milk (solid blue line), dialysed breast milk (dashed red line) and dialysed formula milk (dashed blue line).

**Figure 4:**
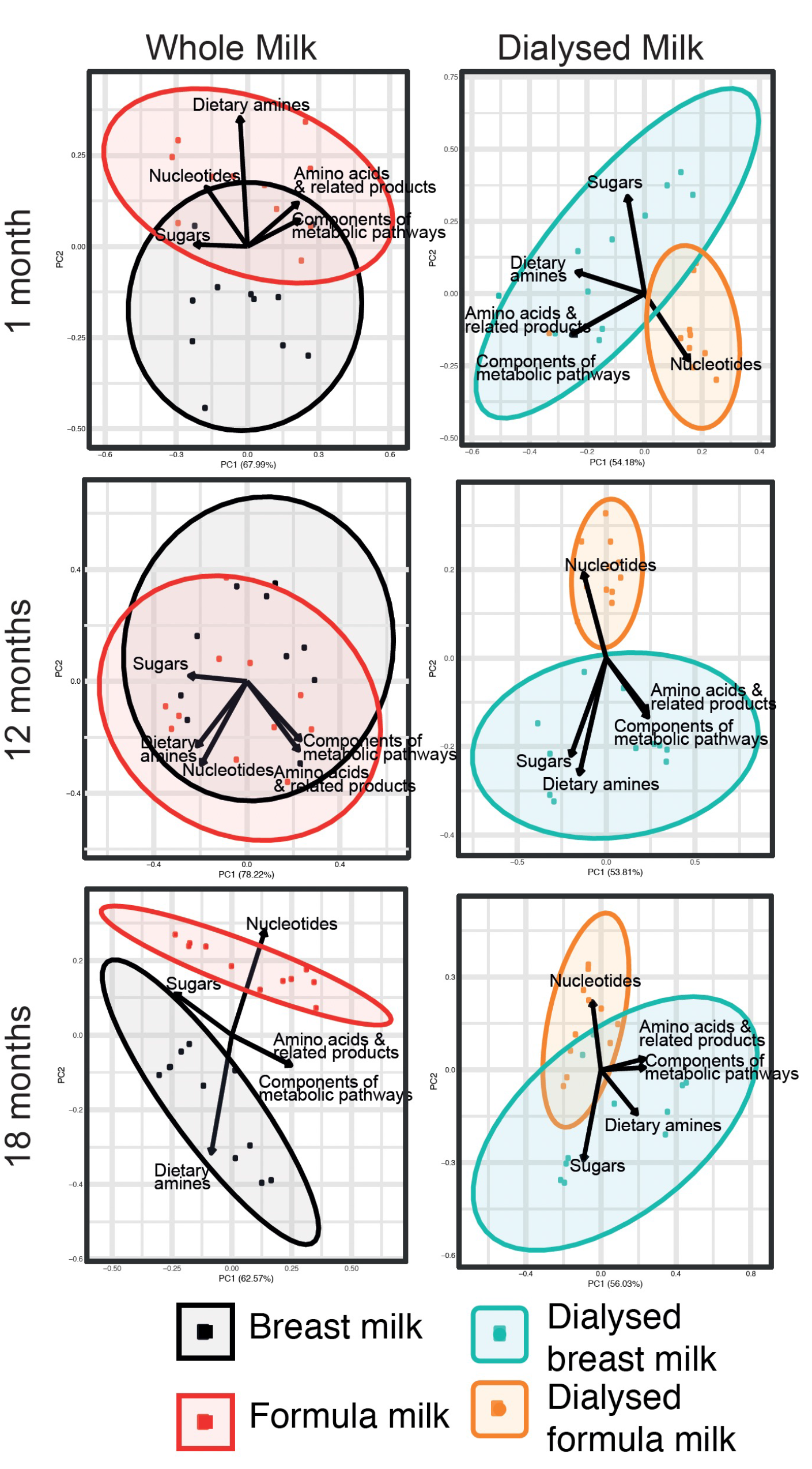
Metabolic profiles during growth in whole and dialysed breast milk and formula. Key metabolites associated with sugars, amino acids & related metabolic products, components of metabolic pathways, nucleotides and dietary amines as identified by ^1^H-NMR.

We also noted there was two distinct lipidomic phases observed in breast milk conditions; the first phase at the start of the experiment and the second phase between 5-8 hours. indicating a biphasic uptake of lipids by the microbiota. In both dialysed milk conditions, lipid concentrations were markedly reduced from the onset. We found that throughout the experiment there was a slight increase of TG46:1 levels in the dialysed breast milk compared to dialysed formula milk (Fig 3B, S5 and S6). This slight increase in lipid concentration over time in dialysed breast milk (but not present in the dialysed formula milk) suggests that the larger lipid components retained after dialysis, (although fewer in number) may still be utilised by the microbiota.

### Metabolomics

To further investigate how the faecal microbiota from each infant responded to the different milk diets we performed ^1^H NMR on all time points. Across all infants, we observed a consistent decrease in sugar compounds over time in both whole milk experimental conditions, suggesting that readily accessible sugars are likely key drivers of microbial growth in these conditions (Fig S7, Table S4). A similar trend, although to a significantly lesser extent, was observed when examining the absolute concentrations of metabolites in dialysed milk conditions (Table S4). Notably, sugars such as fucose were only detected when the microbiota was grown in breast milk or formula, indicating that these sugars likely originate from the diet, and are essential components for microbial metabolism. Additionally, key metabolites produced during microbial fermentation, such as lactate and short chain fatty acids (SCFAs), including acetate, butyrate and propionate, were detected in all three infants during growth in breast milk and formula (Fig S7). Over the course of the experiment, we observed that sugar and energy-producing metabolites decreased, while fermentation end-products like ethanol and acetate increased. These opposing trends indicate active microbial growth and metabolism across all conditions.

When analysing the influence of specific metabolic drivers on microbial composition in each infant, clear differences were observed between breast or formula milk conditions (Fig 4). Metabolites were grouped into categories, including sugars, nucleotides, dietary amines, amino acids, and other metabolic pathway components. In the one-month and 12-month-old infants, shifts in microbial composition were strongly driven by sugars and energy-producing metabolites (Fig 4). In contrast, in the more diverse microbiota of the 18-month-old, dietary amines and nucleotides appeared to play a greater role in shaping the microbiota. Regardless of age, the metabolic profile in dialysed formula conditions was significantly reduced compared to undialysed conditions and was marked by increased proportions of amino acids (Fig 4 and S7). We propose that this observation may reflect microbial cell death, as amino acids can be released from dying cells. Additionally, the concentrations of nucleotides was the main driver for differences in the microbiota between dialysed breast and dialysed formula milk across all infants, which could be released from dying cells and functioning as a compound to sustain microbial communities when other nutrients are limited (Fig 4).

### Diet-induced changes in metabolic pathways

It is well established that even small dietary interventions can influence gut microbiota composition and function. To further explore these effects, we identified significant differences in KEGG-enriched metabolic pathways based on our ^1^H-NMR data (significance threshold 0.000001 ≥ 0.001) (Fig 5). Overall, the youngest infant (1-month-old) had the largest number of significantly KEGG-enriched metabolic pathways, many of which were related to carbohydrate metabolism. These included pathways involved in gluconeogenesis, lactose synthesis, pyruvate metabolism and amino sugar metabolism; and the metabolism of key amino acids such as arginine, proline, aspartate and tyrosine. In addition, pathways involved in the degradation of branched-chain amino-acids (valine, leucine and isoleucine) were also significantly enriched (Fig 5). Surprisingly, we observed a similar pattern, albeit less significant, of pathways upregulated when the one-month-old faeces was grown in dialysed breast milk (Fig 5).

**Figure 5:**
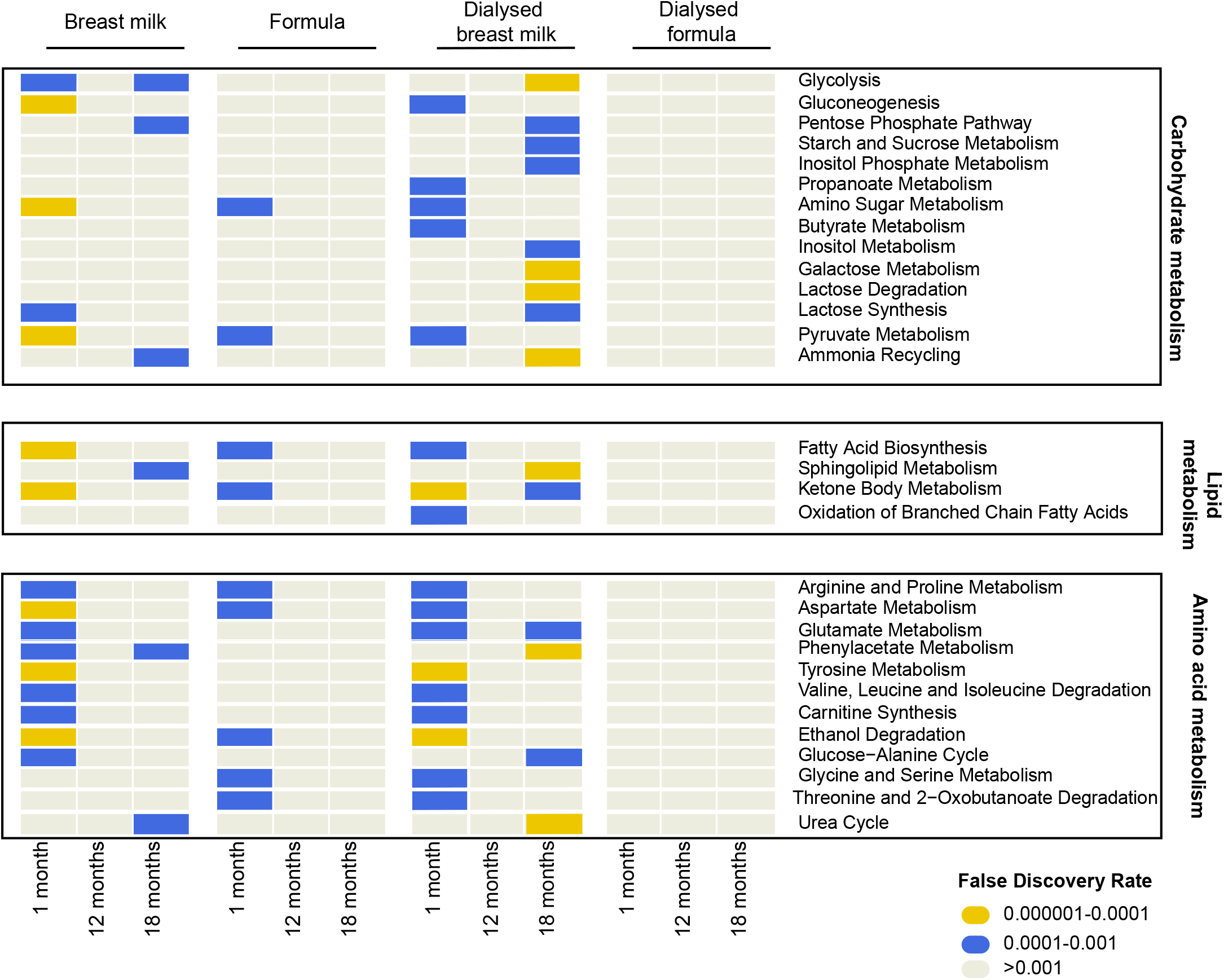
Enriched metabolic pathways identified in carbohydrate, lipid and amino acid metabolism from each infant when grown with whole and dialysed breast milk and formula. KEGG-enriched pathway assignment done using Metaboanalyst and ^1^H-NMR data.

In line with our lipidomic analysis, breast milk conditions also showed enrichment in key lipid-related metabolic pathways, significantly more so than when formula milk was used (Fig 5). These pathways included fatty acid biosynthesis and ketone body metabolism, suggesting that the infant microbiota might be utilising breast milk lipid micelles and associated proteins as an energy source for growth (Fig 5). Notably, lipid metabolism pathways were enriched only in the one-month-old faecal samples when grown in formula milk, and it was the only age group to show any significantly enriched KEGG-pathways under formula conditions. This result suggests that the youngest infant’s microbiota likely retains high plasticity in function, thereby allowing it to rapidly adapt to new environments and metabolites available. This adaptability could also be a reflection the infant’s early transition to a formula diet, where the microbiota had already begun to adapt to the new nutritional environment (Table S1 and S5).

## Discussion

Maternal breast milk is universally recognised as the optimal diet for infants aged 0-6 months (UNCIEF-WHO, 2018), offering essential nutrients and bioactive components that promote healthy growth and development. However, some parents may be unable or choose not to breastfeed and opt for formula milk instead, which while an excellent source of nutrition, cannot fully replicate the unique components of breast milk such as HMOs and maternal-derived immune factors. Additionally, formula feeding does not typically result in the characteristic infant gut microbiome profile seen in exclusively breast-fed infants, where *Bifidobacterium* dominates. What remains unclear is how rapidly the infant microbiome can adapt to different milk diets, which dietary components drive this change, and whether the microbiome retains a form of ‘microbial memory’ of past milk exposures? To address these gaps, we investigated how different milk compositions affect the faecal microbiota of three healthy full-term infants at different developmental stages: 1 month, 12 months, and 18 months. These infants had characteristic age-related microbiota profiles, with *Bifidobacterium* dominating, and increasing taxonomic diversity in line with age of the infant. As many infants receive formula milk to supplement their diet (either partial or full), we examined how these two milk types, under static conditions, could differentially shift the gut microbiota over 48h.

Contrary to our expectations, we found that regardless of infant age, the bacterial composition became less diverse and reverted to a simpler composition dominated by *Bifidobacterium* when exposed to breast milk (Fig 2A-B). This was particularly notable in the older infants, whose initially more diverse microbiota profiles became simplified in the presence of breast milk. *Bifidobacterium* that represented a lower abundance of the total microbiome of the two older infants (Fig S1), bloomed to dominate the microbiota when grown with breast milk. This suggests that the early life microbiota can exhibit a form of ‘microbial memory’ from earlier dietary exposures (Letourneau et al., 2022, Zhang et al., 2023). Such microbial memory has been reported in other contexts, including light-induced memory patterns in microbial communities (Larkin et al., 2018; Yang et al., 2020). Our data support the concept to nutrient-driven microbial memory in the infant gut, where bacteria are not lost when breast milk is removed, but instead persist at low abundance and can bloom again when prebiotics (i.e. HMOs) and reintroduced (Letourneau et al., 2022; Zhang et al., 2023).

Strikingly, the microbiota of the youngest infant (1-month-old) was highly plastic and stochastic across all experimental conditions, especially in formula milk. This is likely due to the high dynamic nature of the microbiome during early life stages, which drives rapid responses to dietary changes. However, this plasticity also likely reflects the developmental immaturity of the infant’s microbiome, as it undergoes constant changes and struggles to establish stability over the time, unlike the older two infants. In particular, the microbiota of the 18-month-old infants was very diversified and yet more stable, and we observed tightly clustered alpha and beta diversity metrics over time across all diet conditions examined. These results suggest that by 18 months, the gut microbiota has achieved a degree of stability, making it more readily resilient to small diet changes.

To investigate the effects of large macromolecules in milk, we dialysed both breast and formula milk to remove smaller sugars and molecules that would typically be absorbed in the upper small intestine, leaving behind larger proteins and fat globules. Although the dialysis functioned to mimic what the microbiome is exposed to in the colon, we did not add lipases and bile acids, which would normally contribute to lipid metabolism in the gut. However, we found that bacterial composition varied depending on the remaining components in each diet. It is known that breast milk contains fat globules and lipids that are of varying sizes and structures, which likely influence which bacteria can use these lipids for energy. In contrast, formula milk contains more uniform lipid emulsions (Furse and Koulman, 2020), which may limit the diversity of bacteria that can use them. Lipids are a key energy source for the developing microbiota, and the variability in breast milk lipidomic profiles may be a critical component in promoting a healthy infant microbiota dominated by *Bifidobacterium*. Interestingly, independent of the infants age – the gut microbiota of breast-fed infants reverted to a simplistic microbial community when re-exposed to breast milk alone, demonstrating a form of microbial memory. In younger infants, we observed that the infant gut microbiota had high plasticity and responded rapidly (within hours) to dietary changes. This was supported by 16S rRNA amplicon analysis and the identification of enriched KEGG-assigned pathways, which showed shifts in metabolic activity in response to different milk types.

The results of this study are based on a single sample from each infant, grown in a static culture rather than a continuous culture system. Although we performed in-depth lipidomic and metabolic profiling of the infants’ microbiome response to milk diets, future studies should ideally include a larger number of infant samples and incorporate shotgun metagenomic sequencing for more strain level insights. Despite these limitations, our findings have implications for understanding how early life diet shapes the gut microbiota and how dietary interventions can be used to support healthy microbial development. The rapid adaptability of the infant microbiome in early life and its ability to revert to simpler states when exposed to breast milk suggest that developing targeted bacterial and milk-based supplements could promote optimal microbiome development during critical early life windows. Further research is needed to explore the potential of these interventions in promoting long-term health outcomes in infants.

## Acknowledgements

We would like to give a special mention to the mothers that donated expressed breast milk for this study, as well as to the families that collected and donated their infant’s stool.

## References

Alcon-Giner C, Caim S, Mitra S, Ketskemety J, Wegmann U, Wain J, et al. (2017) Optimisation of 16S rRNA gut microbiota profiling of extremely low birth weight infants. BMC Genomics. 18(1):841.

Altschul SF, Madden TL, Schäffer AA, Zhang J, Zhang Z, Miller W, et al. (1997 (Gapped BLAST and PSI-BLAST: A new generation of protein database search programs. Nucleic Acids Res. 25(17):3389–402.

Bäckhed F, Roswall J, Peng Y, Feng Q, Jia H, Kovatcheva-Datchary P, et al. (2015) Dynamics and stabilization of the human gut microbiome during the first year of life. Cell Host Microbe. 17:690–703.

Björkstén, B., Sepp, E., Julge, K., Voor, T., & Mikelsaar, M. (2001). Allergy development and the intestinal microflora during the first year of life. Journal Of Allergy And Clinical Immunology, 108(4), 516–520. doi: 10.1067/mai.2001.118130

Ding T, Xu M, Li Y. (2022) An Overlooked Prebiotic: Beneficial Effect of Dietary Nucleotide Supplementation on Gut Microbiota and Metabolites in Senescence-Accelerated Mouse Prone-8 Mice. Front Nutr. Mar 24;9:820799. doi: 10.3389/fnut.2022.820799. PMID: 35399683; PMCID: PMC8988891.

Elison, E., Vigsnaes, L., Rindom Krogsgaard, L., Rasmussen, J., Sørensen, N., & McConnell, B. et al. (2016). Oral supplementation of healthy adults with 2′-O-fucosyllactose and lacto-N-neotetraose is well tolerated and shifts the intestinal microbiota. British Journal Of Nutrition, 116(8), 1356–1368. doi: 10.1017/s0007114516003354

Ferretti P, Pasolli E, Tett A, Asnicar F, Gorfer V, Fedi S, et al. Mother-to-infant microbial transmission from different body sites shapes the developing infant gut microbiome. Cell Host Microbe. 2018;24:133–45.e5

Furse S, Koulman A. The Lipid and Glyceride Profiles of Infant Formula Differ by Manufacturer, Region and Date Sold. Nutrients. 2019 May 20;11(5):1122. doi: 10.3390/nu11051122. PMID: 31137537; PMCID: PMC6567151.

Gratton J et al. An optimized sample handling strategy for metabolic profiling of human feces. Anal. Chem. 2016, 88, 4661–4668.

Huson D, Mitra S, Ruscheweyh H. Integrative analysis of environmental sequences using MEGAN4. Genome Res. 2011;21(9):1552–60.

Kim SY, Yi DY. Components of human breast milk: from macronutrient to microbiome and microRNA. Clin Exp Pediatr. 2020 Aug;63(8):301–309. doi: 10.3345/cep.2020.00059. Epub 2020 Mar 23. PMID: 32252145; PMCID: PMC7402982.

Koenig JE, Spor A, Scalfone N, Fricker AD, Stombaugh J, Knight R, Angenent LT, and Ley RE (2011). Succession of microbial consortia in the developing infant gut microbiome. Proc. Natl. Acad. Sci. U. S. A 108 Suppl 1, 4578–4585.

Larkin, J., Zhai, X., Kikuchi, K., Redford, S., Prindle, A., & Liu, J. et al. (2018). Signal Percolation within a Bacterial Community. Cell Systems, 7(2), 137-145.e3. doi: 10.1016/j.cels.2018.06.005

Laursen, M., Bahl, M., Michaelsen, K., & Licht, T. (2017). First Foods and Gut Microbes. Frontiers In Microbiology, 8. doi: 10.3389/fmicb.2017.00356

Laursen, M., Sakanaka, M., von Burg, N., Mörbe, U., Andersen, D., & Moll, J. et al. (2021). Bifidobacterium species associated with breastfeeding produce aromatic lactic acids in the infant gut. Nature Microbiology, 6(11), 1367–1382. doi: 10.1038/s41564-021-00970-4

Letourneau J, Holmes ZC, Dallow EP, Durand HK, Jiang S, Carrion VM, Gupta SK, Mincey AC, Muehlbauer MJ, Bain JR, David LA. 2022. Ecological memory of prior nutrient exposure in the human gut microbiome. ISME J 16(11):2479–2490. doi: 10.1038/s41396-022-01292-x.

Ma, J., Li, Z., Zhang, W., Zhang, C., Zhang, Y., & Mei, H. et al. (2020). Comparison of gut microbiota in exclusively breast-fed and formula-fed babies: a study of 91 term infants. Scientific Reports, 10(1). doi: 10.1038/s41598-020-72635-x

Nicholson JK, et al. 750 MHz 1H and 1H-13C NMR spectroscopy of human blood plasma. Anal Chem 1995, 67, 793–811.

O’Neill, I., Schofield, Z., & Hall, L. (2017). Exploring the role of the microbiota member Bifidobacterium in modulating immune-linked diseases. Emerging Topics In Life Sciences, 1(4), 333–349. doi: 10.1042/etls20170058

Psychogios N, et al, The human serum metabolome. PloS One, 2011, 6:e16957

Quast C, Pruesse E, Yilmaz P, Gerken J, Schweer T, Yarza P, et al. The SILVA ribosomal RNA gene database project: Improved data processing and web-based tools. Nucleic Acids Res. 2013;41(D1):590–6.

Rollins, N.C.; Bhandari, N.; Hajeebhoy, N.; Horton, S.; Lutter, C.K.; Martines, J.C.; Piwoz, E.G.; Richter, L.M.; Victora, C.G. Why invest, and what it will take to improve breastfeeding practices? Lancet 2016, 387, 491–504.

Stewart, C., Ajami, N., O’Brien, J., Hutchinson, D., Smith, D., & Wong, M. et al. (2018). Temporal development of the gut microbiome in early childhood from the TEDDY study. Nature, 562(7728), 583–588. doi: 10.1038/s41586-018-0617-x

Sun, S., Luo, L., Liang, W., Yin, Q., Guo, J., & Rush, A. et al. (2020). Bifidobacterium alters the gut microbiota and modulates the functional metabolism of T regulatory cells in the context of immune checkpoint blockade. Proceedings Of The National Academy Of Sciences, 117(44), 27509–27515. doi: 10.1073/pnas.1921223117

UNCIEF-WHO. 2018 Capture the Moment-Early initiation of breastfeeding: The best start for every newborn. <http://www.who.int/nutrition/publications/infantfeeding/capture-moment-early-initiation-bf/en/2018>; 41 New York: UNICEF.

Victora, C.G.; Bahl, R.; Barros, A.J.D.; França, G.V.A.; Horton, S.; Krasevec, J.; Murch, S.; Sankar, M.J.; Walker, N.; Rollins, N.C. Breastfeeding in the 21st century: Epidemiology, mechanisms, and lifelong effect. Lancet 2016, 387, 475–490.

Weljie AM, et al. Targeted profiling: quantitative analysis of 1H NMR metabolomics data. Anal Chem 2006, 78, 4430–4442.

Yang, C., Bialecka-Fornal, M., Weatherwax, C., Larkin, J., Prindle, A., & Liu, J. et al. (2020). Encoding Membrane-Potential-Based Memory within a Microbial Community. Cell Systems, 10(5), 417-423.e3. doi: 10.1016/j.cels.2020.04.002

Yassour M, Jason E, Hogstrom LJ, Arthur TD, Tripathi S, Siljander H, et al. Strain-level analysis of mother-to-child bacterial transmission during the first few months of life. Cell Host Microbe. 2018;24:146–54.e4.

Yatsunenko T, Rey FE, Manary MJ, Trehan I, Dominguez-Bello MG, Contreras M, et al. 2012 Human gut microbiome viewed across age and geography. Nature. 486:222–7.

Zhang C, Kong Y, Xiang Q, Ma Y, Guo Q. 2023. Bacterial memory in antibiotic resistance evolution and nanotechnology in evolutionary biology. iScience. 26(8):107433. doi: 10.1016/j.isci.2023.107433. PMID: 37575196; PMCID: PMC10415926.

