## Supplementary Figures for "Early life milk diets shape infant gut microbiota: evidence of microbial plasticity in response to breast and formula milk"

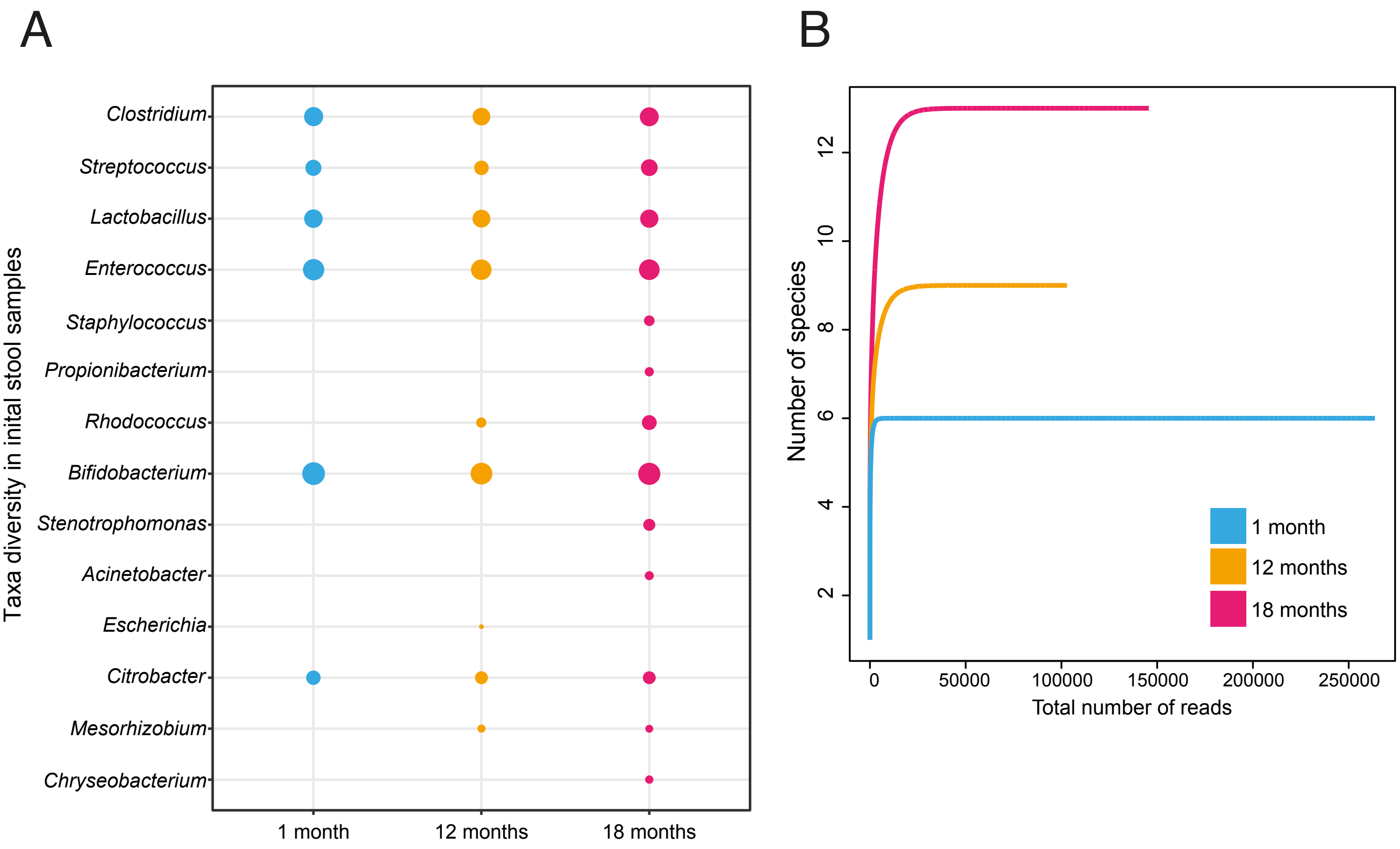


**Supplementary figure 1:** (A) 16S rRNA profile of initial faecal samples for the three infants used in this study. Charts show bacteria diversity in original faecal sample that were subsequently used for growth in breast milk, formula, dialysed breast milk and dialysed formula. Bubble size reflects total number of reads assigned to bacterial genus from 16S amplicon sequencing (B) Refraction curves for 16S amplicon, MiSeq. Graph show the total number of species (y-axis) over the total number of reads (x-axis).

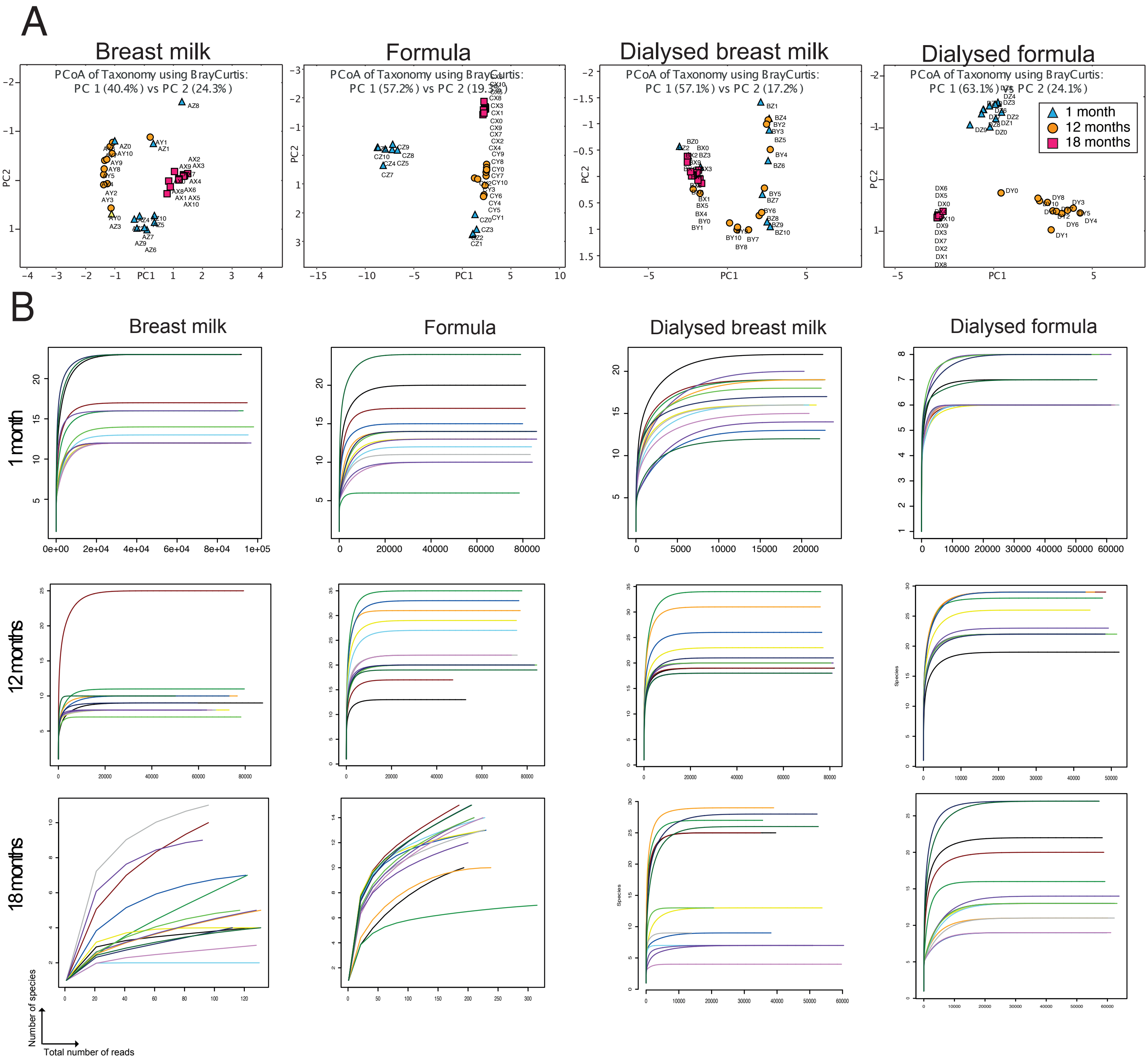


**Supplementary figure 2:** Changes in bacterial composition over time in each specified condition for all three infants. (A) PCA plots of overall changes in bacterial taxonomy. Each point represents a single time point for each infant. (B) Refraction curves for the 16S rRNA amplicon data, each line represents a single time point for each experimental condition.

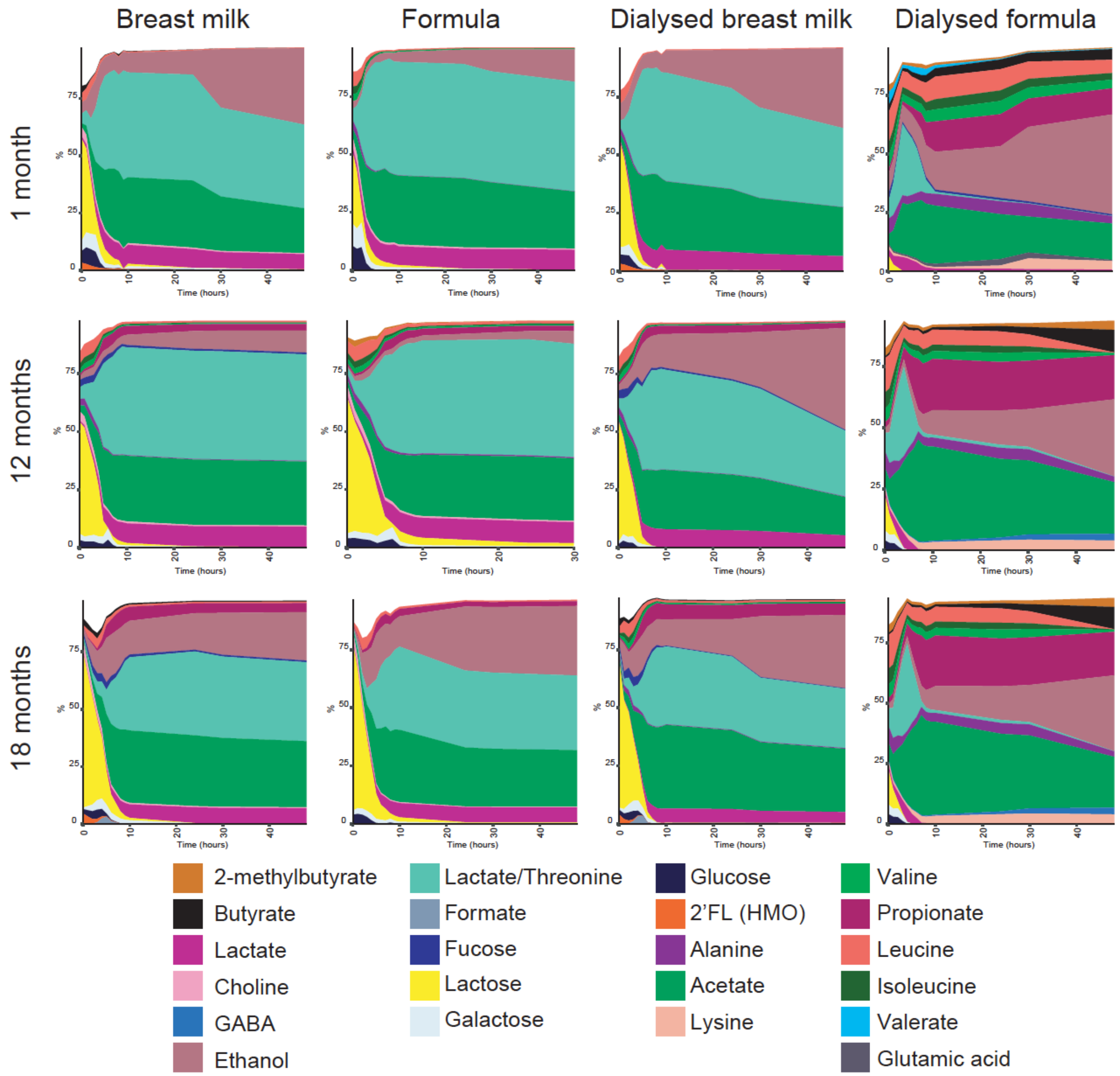


**Supplementary figure 4:** Proportional changes in top 24 metabolites identified for each infant grown in breast milk, formula milk, dialysed breast milk and dialysed formula milk.

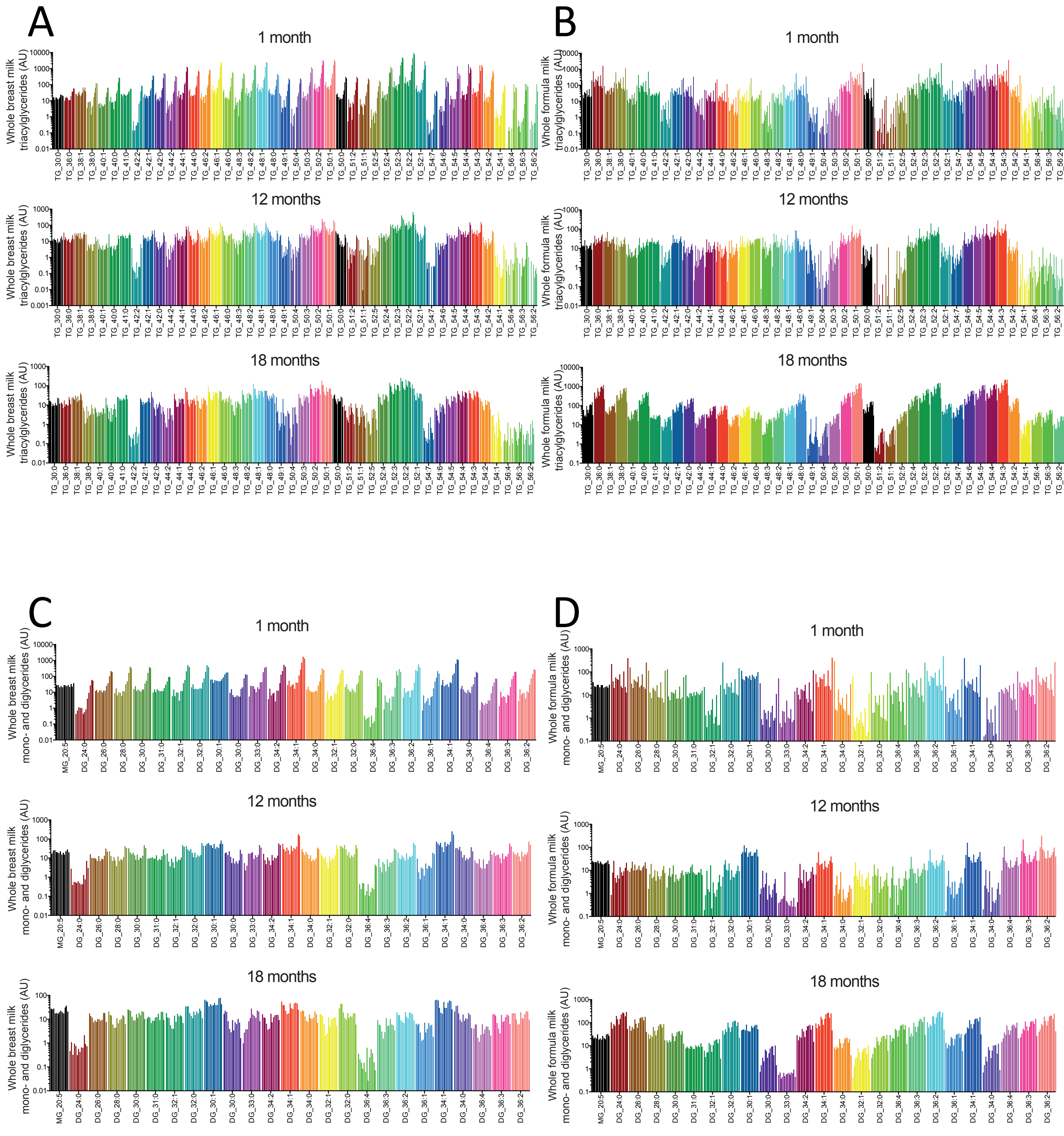


**Supplementary figure 4:** Concentration of triacyglycerides (A&B) and monoglyceride/diacylglycerides (C&D) over 48h in whole breast and formula milk. Each bar represents an individual measurement; x-axis represents change in time (0h-48h) for each lipid.

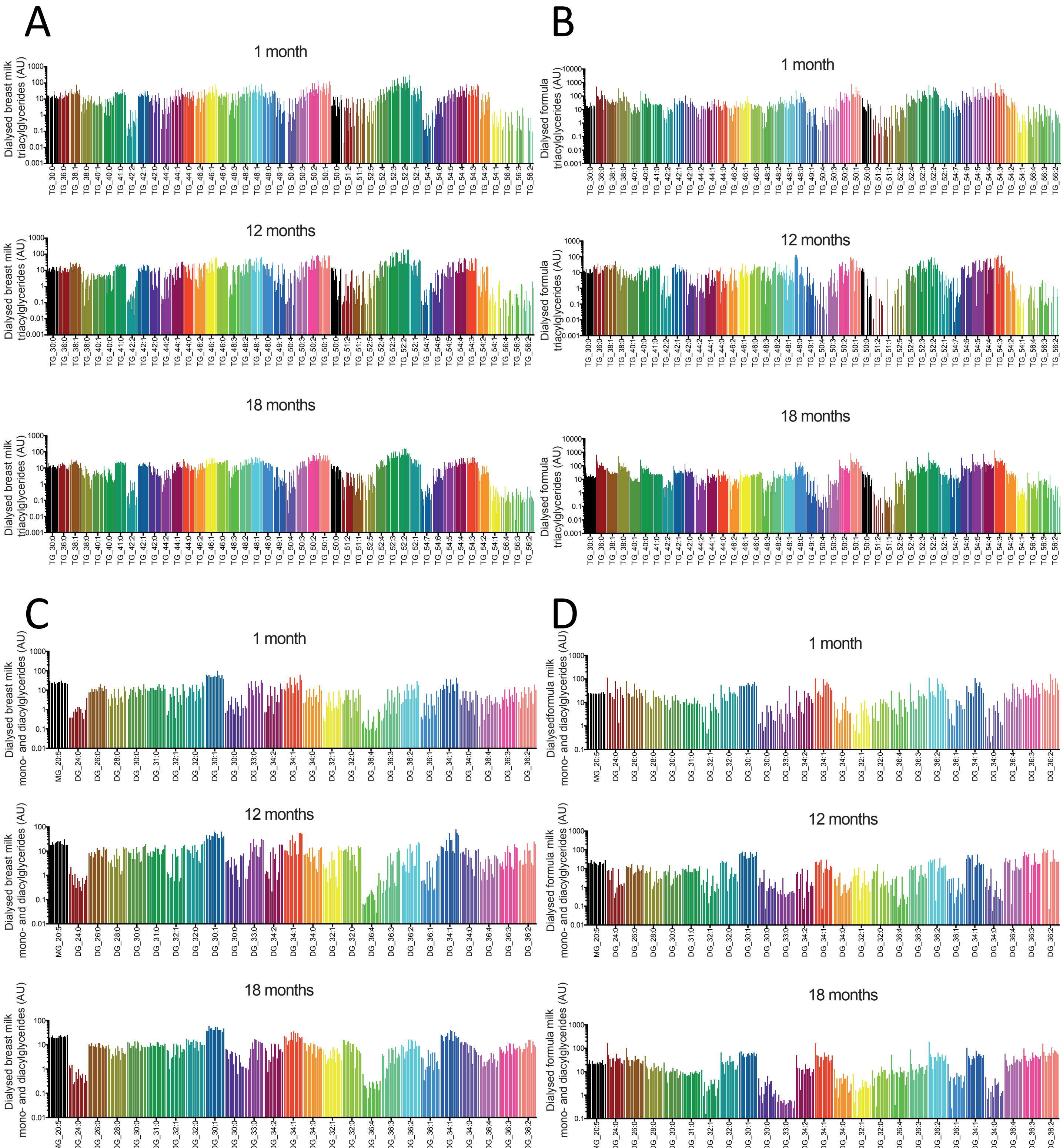


**Supplementary figure 5:** Concentration of triacyglycerides (A&B) and monoglyceride/diacylglycerides (C&D) detected over 48h in dialysed breast and formula milk. Each bar represents an individual measurement; x-axis represents change in time (0h-48h) for each lipid.

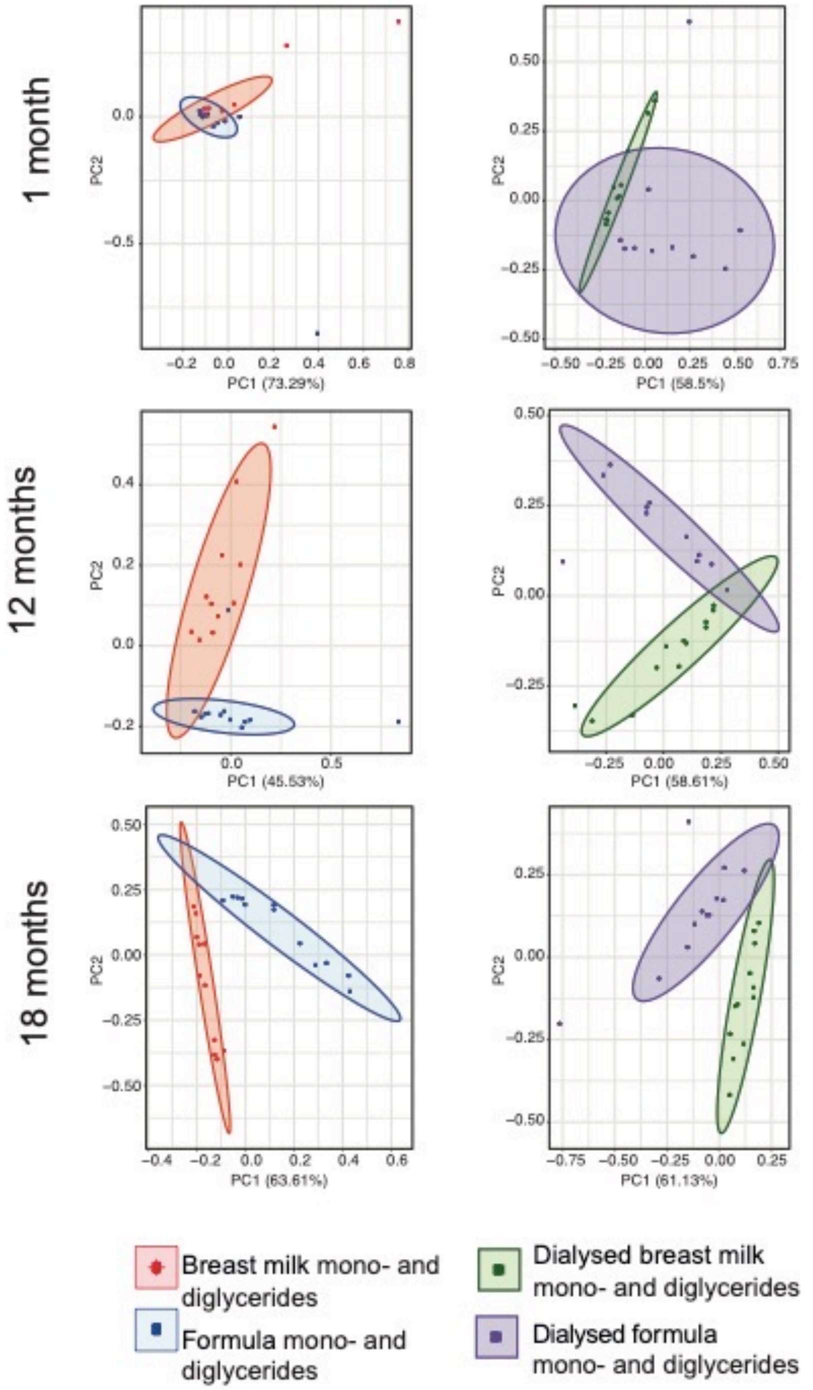


**Supplementary figure 6:** PCA plot for changes in milk monoglyceride and diglycerides for each infant over time (0-24h).

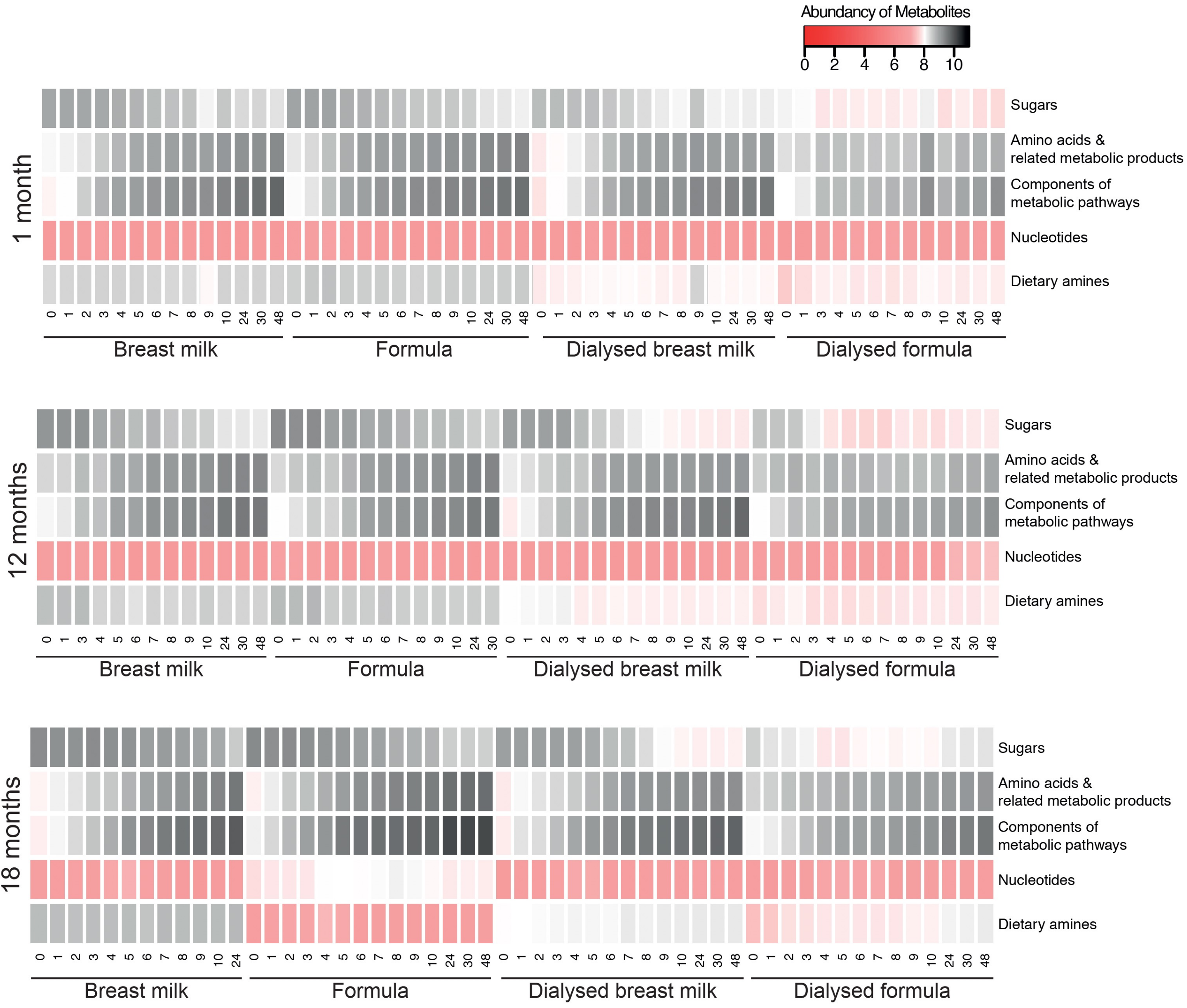


**Supplementary figure 7:** Concentration profile of key metabolite groups identified by $^{1}$H-NMR detected at each time point over 48h in whole breast milk, formula milk, dialysed breast milk and dialysed formula milk for each infant as labelled. Each bar represents an individual measurement; x-axis represents change in time (0h-48h) for each metabolite.
